# Quantifying the impact of public omics data

**DOI:** 10.1101/282517

**Authors:** Yasset Perez-Riverol, Andrey Zorin, Gaurhari Dass, Mihai Glont, Juan Antonio Vizcaíno, Andrew F. Jarnuczak, Robert Petryszak, Peipei Ping, Henning Hermjakob

**Affiliations:** European Molecular Biology Laboratory, EMBL-European Bioinformatics Institute (EMBL-EBI), Hinxton, Cambridge, UK; Department of Physiology and Department of Medicine, Division of Cardiology, David Geffen School of Medicine at UCLA, University of California, Los Angeles, Los Angeles, California, USA; Department of Medicine, Division of Cardiology, David Geffen School of Medicine at UCLA, University of California, Los Angeles, Los Angeles, California, USA; State Key Laboratory of Proteomics, Beijing Proteome Research Center, Beijing Institute of Lifeomics; National Center for Protein Sciences (The PHOENIX Center, Beijing), Beijing 102206, China

**Author notes:** Corresponding author: Dr. Yasset Perez-Riverol, European Molecular Biology Laboratory, European Bioinformatics Institute (EMBLEBI), Wellcome Trust Genome Campus, Hinxton, Cambridge, CB10 1SD, UK.

## Abstract

The amount of omics data in the public domain is increasing every year [1, 2]. Public availability of datasets is growing in all disciplines, because it is considered to be a good scientific practice (e.g. to enable reproducibility), and/or it is mandated by funding agencies, scientific journals, etc. Science is now a data intensive discipline and therefore, new and innovative ways for data management, data sharing, and for discovering novel datasets are increasingly required [3, 4]. However, as data volumes grow, quantifying its impact becomes more and more important. In this context, the FAIR (Findable, Accessible, Interoperable, Reusable) principles have been developed to promote good scientific practises for scientific data and data resources [5]. In fact, recently, several resources [1, 2, 6] have been created to facilitate the Findability (F) and Accessibility (A) of biomedical datasets. These principles put a specific emphasis on enhancing the ability of both individuals and software to discover and re-use digital objects in an automated fashion throughout their entire life cycle [5]. While data resources typically assign an equal relevance to all datasets (e.g. as results of a query), the usage patterns of the data can vary enormously, similarly to other “research products” such as publications. How do we know which datasets are getting more attention? More generally, how can we *quantify* the scientific impact of datasets?

---

Recently, several authors [7-9] and resources [10] pointed out the importance of evaluating the impact of each research product, including datasets. Reporting scientific impact is indeed increasingly relevant for individuals, but also reporting aggregated information has become essential in the case of research groups, scientific consortia, institutions or for public data resources among others, in order to assess the level of importance, excellence and relevance of their work. This is a key piece of information for funding agencies, which is used routinely to prioritize the projects and resources they fund. However, most of the efforts nowadays focus on the evaluation and quantification of the impact of publications as the main artefact. For instance, in 2013, the “altmetrics” team proposed a set of ‘alternative’ metrics to trace research products with special focus on publications [10]. Specific tools and services have been built since to aggregate “altmetrics”, including for instance counts of mentions of a given publication in blog posts, tweets and articles in mainstream media. The “altmetrics” score is widely used by the research community nowadays (e.g. by multiple scientific journals), as a measure of scientific impact. In addition to the traditional journals, a relatively recent trend has been the introduction of pre-print servers in the biological/biomedical field (e.g. bioRxiv), supporting the pre-publication of manuscripts which are usually under review in traditional journals. Pre-prints are increasingly tracked and recognised as research products (e.g. by funding agencies such as The Wellcome Trust). However, adequate tracking and recognition of datasets has been limited so far for multiple reasons: i) the relatively low number of publications citing datasets instead of their corresponding publications; ii) the lack of services that store and index datasets from heterogeneous origins; and iii) the absence of widely-used metrics that enable the quantification of their impact. Some attempts have been made to improve the situation, by introducing DOIs (Data Object Identifiers) directly associated to datasets [11].

In 2016, we released the first version of the Omics Discovery Index (OmicsDI – www.omicsdi.org) as a light-weight system to aggregate datasets across multiple public omics data resources. OmicsDI integrates genomics, transcriptomics, proteomics, metabolomics, and multi-omics datasets, as well as computational models of biological processes [1]. The OmicsDI web interface and Application Programming Interface (API) provide different views and search capabilities on the indexed datasets. Datasets can be searched and filtered based on different types of technical and biological annotations (e.g., species, tissues, diseases, etc.), year of publication, and the original data repository where they are stored, among others. At the time of writing (March 2018), OmicsDI stores just over 100,000 datasets from 16 different public data resources (www.omicsdi.org/database). The split per omics technology is as follows: transcriptomics (68,648 datasets), genomics (12,274), proteomics (9,352), metabolomics (1,679), multi-omics (4,057) and biological models (8,418). Here, we propose a set of novel metrics to quantify the impact of biomedical datasets. A complete framework (now integrated in OmicsDI) has been implemented in order to provide and evaluate those metrics. Finally, we propose a set of recommendations for authors, journals, and data resources to promote an optimal quantification of the impact of datasets.

## Online Methods

In contrast to publications, where the impact is mainly measured by the number of citations, we believe the impact of datasets should be quantified using more than one metric. With this in mind, we have devised three metrics that can be used to provide the impact for datasets:

1. **Number of reanalyses/re-use (*Reanalyses*)**: A reanalysis can be generally defined as the complete or partial re-use of an original dataset (A) using a different analysis protocol, and stored either in the same or in another public data resource (B). For example, PeptideAtlas [12] systematically reanalyses public proteomics datasets, mainly from PRIDE [13] and MassIVE (massive.ucsd.edu). The new peptide and protein evidences from these reanalyses have become an invaluable resource, e.g. for the Human Proteome Project [14]. The appropriate and accurate reference to the original datasets in other resources facilitates the reproducibility and traceability of the results and the recognition for the authors that generated the original dataset [15].
2. **Direct citations of dataset identifiers in publications (*Citations*)**: *Citations* represent the number of publications that directly refer to dataset identifiers [16]. Currently, scientists typically cite publications rather than datasets. However, the citation of the dataset by using the corresponding publication does not imply in our view that the authors have re-used the actual data. Actually, it is more common that citations of papers reference results, claims and biological conclusions rather than the actual dataset. In 2013, Piwowar *et al.* proposed the use of **direct citations** of datasets in the literature as a metric [16], based on the counts generated using the EuropePMC API [17]. EuropePMC stores scientific publications (and also pre-prints, if they have been manually added by the authors) that are open access, so the whole text of the manuscript can be used for performing text mining, enabling this functionality. It is important to highlight that authors can submit their own reanalyses of existing datasets to the same (where the original (reanalysed) datasets was made available) or other public data resources. This information can be used to create a “network” of dataset re-usage.
3. **Number of biological claims based on the dataset (*Connections*):** Here, we quantify the number of biological entities which are supported by a given dataset, in various popular biomedical “knowledge-base” databases such as UniProt [18] (protein sequences), IntAct [19] (molecular interaction data) or Reactome [20] (biological pathways) (**Supplementary Note 2**). Most of the biomedical datasets support biological claims (e.g. pathways, interactions, expression profiles, etc.) that are either manually curated, or automatically annotated in relevant knowledge-bases. We believe that calculating impact in this manner is needed, since for instance, if a number of gene expression profiles were supported by one particular dataset, that information would be lost (or at least untraceable) if the dataset were no longer publicly available. All these knowledge-base databases are indexed and queried by OmicsDI using the EBI Search indexing system (www.ebi.ac.uk/ebisearch/)[21](**Supplementary Note 2**).

## Proposed platform to quantify dataset impact

In order to compute the proposed metrics (*Reanalyses, Citations, Connections*), a novel platform has been developed within OmicsDI. Figure 1 shows a schematic representation of the platform. OmicsDI imports the metadata for each dataset from the original providers, using the OmicsDI XML file format (**Fig. 1a**) [1]. In order to ensure that the metrics are accurate, the infrastructure implements a system to remove dataset redundancy (when two different resources store the same original dataset). An automatic pipeline and a manual annotation system enable OmicsDI to remove duplicated datasets with potentially different identifiers (e.g. transcriptomics datasets available in ArrayExpress and Gene Expression Omnibus (GEO)) (**Fig. 1b**). The pipeline (**Fig. 1b**) designates one of the datasets as the canonical representation and annotates the rest of identifiers as additional secondary ones (**Supplementary Note 1**).

**Figure 1:**
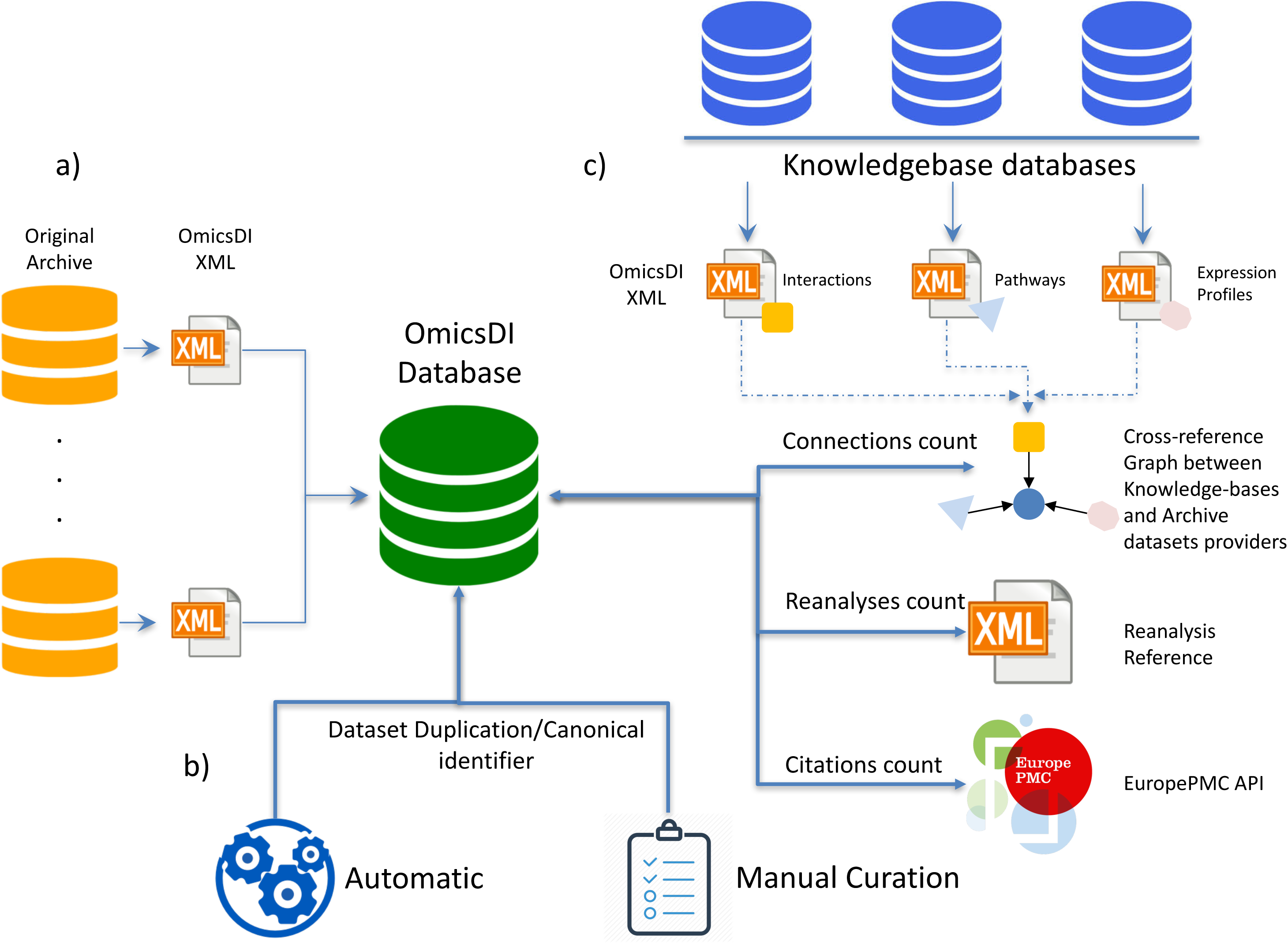
The OmicsDI dataset impact quantification framework allows to compute three metrics (*reanalyses, citations* and *connections*). **a)** OmicsDI datasets are imported from the original public data resources, which are represented in the OmicsDI XML schema. Datasets are imported into a MongoDB database; **b)** the identifier resolution framework allows the detection of datasets that are duplicated across OmicsDI. The final dataset entry contains all the dataset identifiers available in different resources; **c)** *Reanalyses* are counted using the cross-reference system implemented in the OmicsDI XML schema, where a reanalysed dataset can reference the original one. *Citations* are compiled using the EuropePMC API. *Connections* are computed using the biological entities stored in knowledge-bases.

By March 2018, 35 knowledge-bases were used to identify and count the number of biological *connections* (**Supplementary Note 2**). Each tracked biological entity available in the knowledge-bases includes a cross-reference to original datasets (**Fig. 1c**). Then, a software component in OmicsDI navigates this information and computes the *connections* for each of them. The implementation of the *reanalysis* metric is possible because the OmicsDI XML schema (https://github.com/OmicsDI/specifications) provides a mechanism to define when a dataset is based on another one. Finally, the *citations* metric is computed by parallel calls to the EuropePMC API (europepmc.org/RestfulWebService) [17] and the annotation of the final counts for each dataset. All these pipelines and software components can scale to handle thousands of datasets and systematically compute and update the metrics on a weekly basis (github.com/OmicsDI/index-pipeline).

## Dataset Claiming Component

Analogously to services such as Google Scholar and ResearchGate for publications, we have implemented a mechanism that enables researchers to create their own profile in OmicsDI, by claiming their own datasets. Researchers need to log into OmicsDI using their corresponding ORCID account details (**Supplementary Note 3**), and search for relevant datasets using different criteria such as: (i) dataset identifiers, (ii) specific keywords in the title or description of the dataset, or analogous information from the corresponding manuscript where the generation of the dataset is reported, and/or (iii) the author’s name, among others. Then, datasets can be added to an OmicsDI personal profile where it is possible to visualise the impact metrics (*reanalyses, citations* and *connections)*, providing researchers this rich information. The URLs of personal profiles can be shared with anyone in the community. Additionally, as a key point, OmicsDI claimed datasets can be synchronized to the researcher’s own ORCID profile, highlighting datasets there as a research product as well [22] (**Supplementary Note 3**). Although this mechanism is initially aimed at individual researchers (e.g. www.omicsdi.org/profile/Da9hYZOs), research groups, scientific consortia (e.g. www.omicsdi.org/profile/ZEd3mwfF), and research institutions can also create their own OmicsDI profile, facilitating the aggregation, visualisation, tracking and impact assessment for their generated datasets, and the addition to their own OmicsDI profiles. In addition, OmicsDI has implemented a simple visualization component (**Supplementary Note 5**) that allows users to cite the corresponding dataset using the FORCE11 Data Citation Synthesis Group recommendations (http://www.dcc.ac.uk/resources/how-guides/cite-datasets)[23].

## Results

The *reanalyses* metric quantifies how many times one dataset has been re-used (re-analysed) and the result deposited in the same or in another resource. By March 2018, the number of datasets that have been reanalysed at least once is 8,000. The number of reanalyses varies depending on each field and on the existing integration among databases. For example, on average BioModels models are reused 0.8 times, whereas PRIDE (proteomics), MassIVE (proteomics) and ArrayExpress (transcriptomics) datasets are reused 0.12, 0.01 and 0.18 times, respectively.

BioModels MODEL1402200003 (“Genome-scale metabolic modelling of hepatocytes reveals serine deficiency in patients with non-alcoholic fatty liver disease”) [24] is the most re-used “dataset” (the concept of dataset in OmicsDI is extended to biological models), with 6,691 re-uses made available in the same BioModels resource. However, it should be noted that all re-uses are associated with a single publication [25], in which patient data was used to parameterise MODEL1402200003 to 6,691 instances. Frequently, dataset re-use is a hierarchical process, where one dataset is reanalysed subsequently multiple times. **Figure 2a** presents a “reanalysis network” for the model BIOMD0000000055, starting from 2006 (release year) to 2015. A different pattern is illustrated in **Figure 2b**, where BIOMD0000000286 is derived from multiple source models. BioModels curates and annotates for each deposited model, the corresponding model from which it *is derived* (if applicable). **Figure 2c** shows the reanalysis network for three other resources: PRIDE (proteomics), MassIVE (proteomics) and ArrayExpress (transcriptomics). The most reanalysed datasets are PXD000561 (75 times) and PXD000865 (61) corresponding to the two “drafts of the human proteome”. These datasets have supported the annotation of millions of peptides and proteins evidences, enabling the large-scale annotation of the human proteome [14] and have been reanalysed by multiple databases including the proteomics resources PeptideAtlas and GPMDB [26]. In contrast to biological models, the proteomics and transcriptomics fields are still working to define a proper mechanism to report the multiple reanalysis of datasets in a hierarchical manner [11]. Interestingly, the distribution of the elapsed time between the year of publication of the original datasets and publication of the reanalyses shows that most of the datasets are reanalysed within one year after publication (**Fig. 3a**). In fact, three years after the release date, typically the number of reanalyses decrease significantly, making this a metric better suited to measure immediate impact.

**Figure 2:**
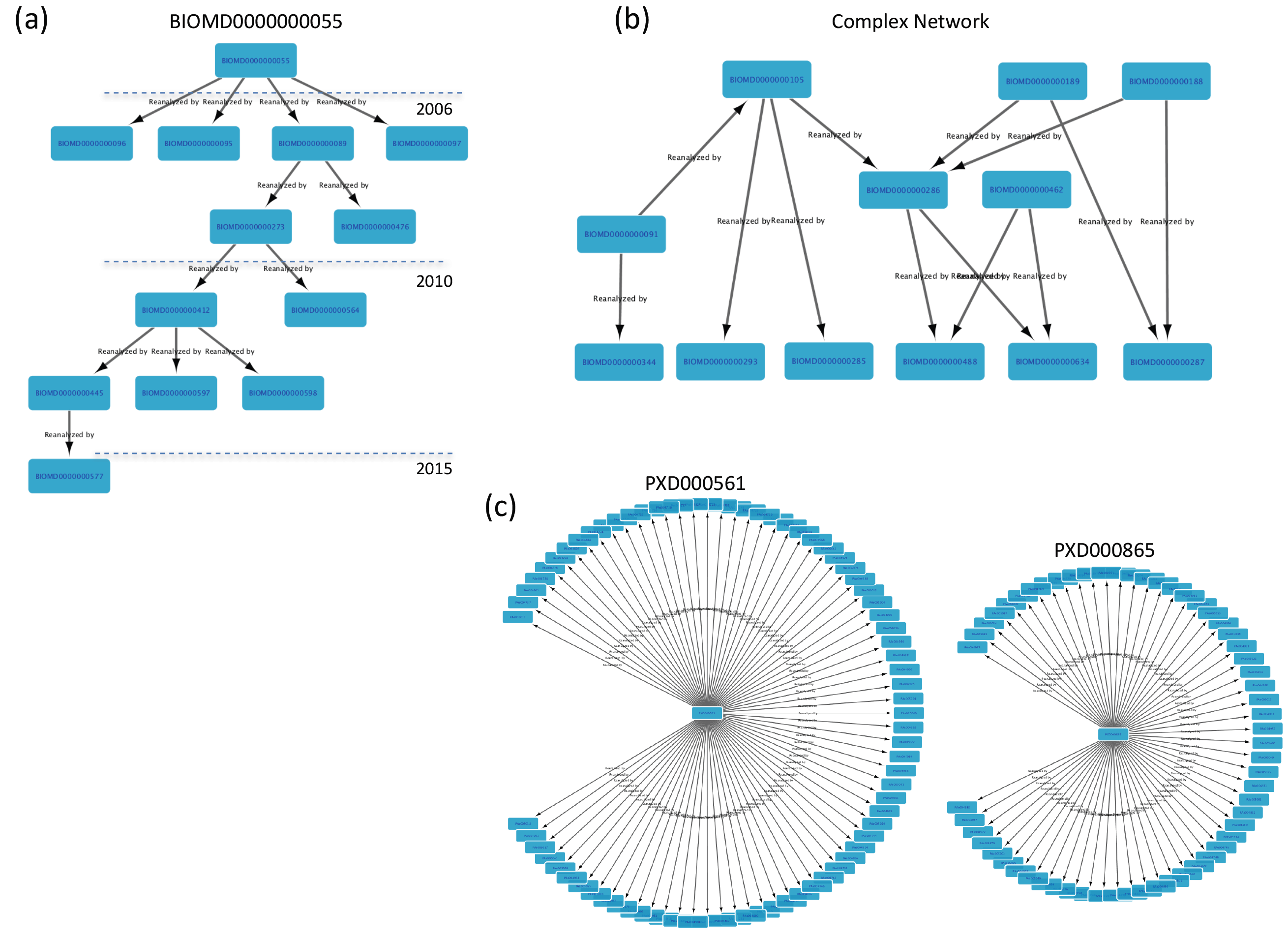
Examples of the reanalysis network for different OmicsDI datasets: **a)** BioModels model BIOMD0000000055; **b)** 12 different BioModels models; **c)** Datasets corresponding to the two drafts of the human proteome (PXD000561 and PXD000865).

**Figure 3:**
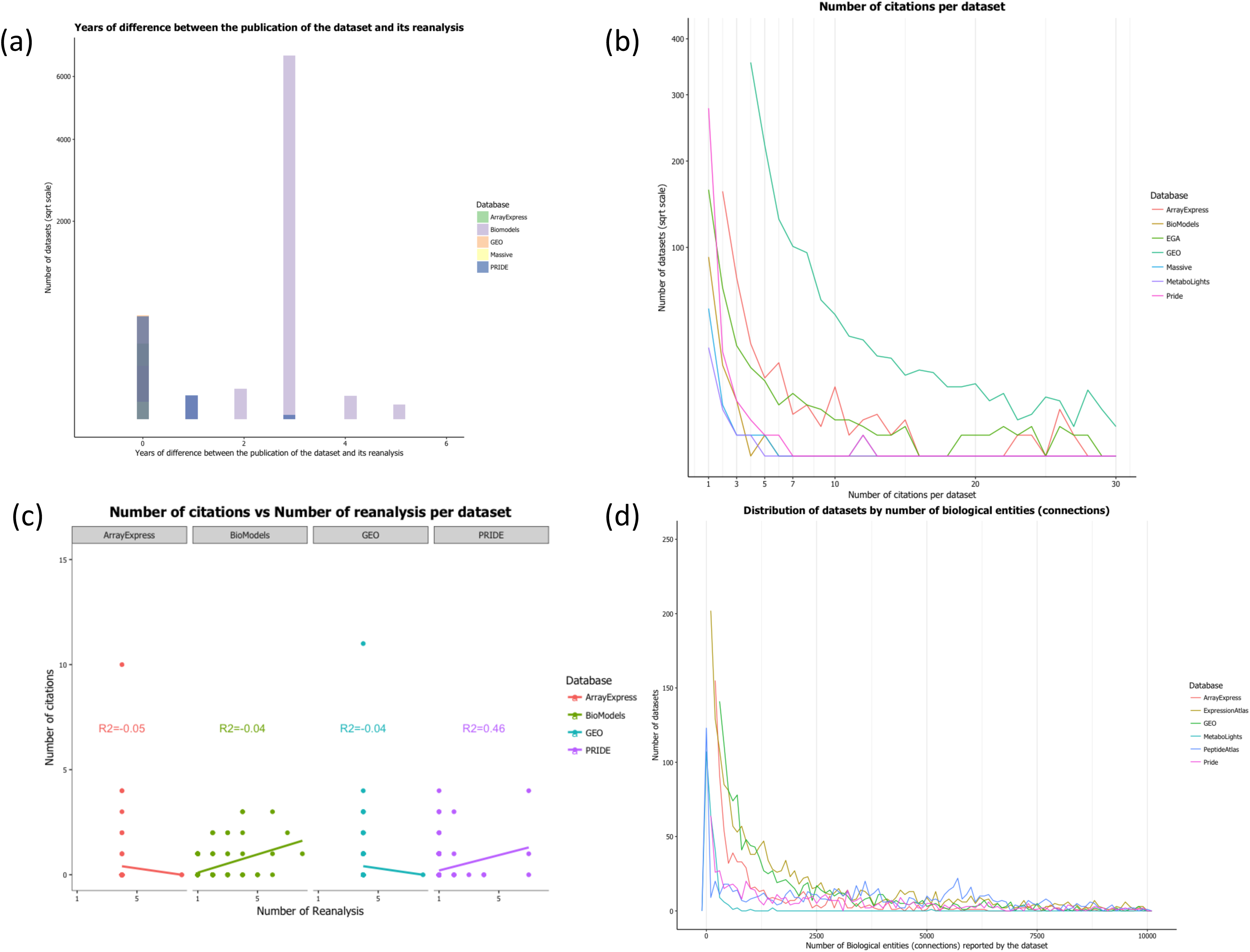
**a)** Elapsed time between the original publication of a dataset and the publication of all its reanalyses; **b)** Distribution of the number of citations per dataset, per OmicsDI data resource; **c)** Correlation between the number of citations per dataset and the number of reanalyses for ArrayExpress, BioModels, GEO and PRIDE datasets; **d)** Distribution of the number of biological connections per dataset.

The second metric is the number of direct *citations* in publications for each dataset as previously suggested [16]. We obtained this information from the EuropePMC API. The number of datasets with at least one citation in EuropePMC is 8,954, including: 6,870 from GEO [27], 982 from ArrayExpress, 343 from EGA (European Genotype Archive, genomics/transcriptomics restricted-access patient data) [28], 316 from PRIDE [13], 118 from BioModels, 60 from MassIVE, 34 from MetaboLights (metabolomics) [29], 3 from PeptideAtlas [12], and 2 from GNPS (metabolomics) [30]. **Figure 3b** shows the distribution of citations per dataset, across seven resources. GEO datasets are the most cited ones, followed by those from ArrayExpress and EGA. The average (and standard deviation of the) number of citations per dataset for each of the five major databases: GEO (transcriptomics), ArrayExpress (transcriptomics), EGA (genomics/transcriptomics), PRIDE (proteomics) and BioModels (models) is 3.4 (9.8), 2.5 (7.2), 4.2 (0.8) and 1.2 (0.66), respectively. Interestingly, the standard deviation indicates that in transcriptomics - GEO (9.8) and ArrayExpress (7.2)some datasets get significantly more attention from the community than others, whereas for proteomics datasets – PRIDE (0.8) – the citation rate is much more homogenous.

The current workflow searches EuropePMC using all the identifiers associated with a given dataset (e.g. a given dataset can be cited in a publication using the ArrayExpress, GEO or BioProject identifiers). For example, the dataset E-GEOD-2034 (www.omicsdi.org/dataset/arrayexpress-repository/E-GEOD-2034) is cited 312 and 28 times using the ArrayExpress (E-GEOD-2034) and GEO (GSE2034) identifiers, respectively. We have correlated the number of dataset citations and reanalyses in four of the major databases (GEO, ArrayExpress, PRIDE and BioModels, **Fig. 3c**). It can be observed that the counts of *citations* in publications and *reanalyses* are not correlated for most resources (GEO, ArrayExpress and BioModels). However, in proteomics (PRIDE) most of the datasets that are re-used by the same or other resources (reanalyses), are also cited by independent researchers in publications (R^2^ = 0.46, p-value = 8.72e-31). This shows that both features are orthogonal, complementary and can be used in combination to get a broader representation of the impact of omics datasets.

Finally, we analysed the number of *connections* that are supported by each dataset. The average (and standard deviation) number of *connections* per dataset for ArrayExpress, ExpressionAtlas, GEO, MetaboLights, PeptideAtlas and PRIDE are 1,253 (31,212), 4,646 (37,842), 466 (6,336), 145 (372), 3,997 (3,477), and 1,556 (13,708); respectively (**Fig. 3d**). The distribution of *connections* per resource shows that many of the datasets in OmicsDI (24%) are connected to at least one biological entity, and most of the distributions follow a similar pattern. For example, dataset E-MTAB-599 (“RNA-seq of mouse DBA/2J x C57BL/6J heart, hippocampus, liver, lung, spleen and thymus”), associated with this publication [31], has 1,710,979 connections, including 1,689,177 genome variants, 21,572 gene values, and 230 other connections, going from sample annotations to nucleotide sequences.

## Discussion

One of the obstacles to achieving a systematic deposition of datasets in public repositories is the lack of a broad scientific reward system, considering other research products in addition to scientific publications [7]. Different studies have demonstrated the need for metrics and frameworks to quantify the impact of deposited datasets in the public domain [7, 16, 23]. Such a system would not only encourage authors to make their data public, but also help funding agencies, biological resources and the scientific community as a whole to focus on the most impactful datasets. In OmicsDI we have implemented a novel platform to quantify the impact of public datasets systematically, by using data from biological data resources (*reanalyses*), literature (*citations*), and knowledge-bases (*connections*). Every metric is updated on a weekly basis and made available through the OmicsDI web interface and API (http://www.omicsdi.org/ws/).

One of the primary findings is that in systems biology (the BioModels database is the representative resource), the deposition of data has enabled systematically generation of new knowledge (biological models), based on previous datasets. For example, the model “Genome-scale metabolic modelling of hepatocytes reveals serine deficiency in patients with non-alcoholic fatty liver disease” (MODEL1402200003) [24] has been used to build more than 6,000 models available in BioModels. Moreover, the results showed that the *reanalyses* metric is crucial to highlight relevant datasets early after the dataset release (**Fig. 2c**). 8,000 datasets (>5% of OmicsDI) have been reanalysed by resources such as PeptideAtlas, GPMDB, or the EBI Expression Atlas, among others. However, it should be noted that the *reanalyses* metric measures only the impact of datasets in the same or other data resources contributing their metadata to OmicsDI, which constitutes a fraction of the total re-use by the scientific community.

To complement the *reanalysis* metric, we counted direct citations of datasets in scientific publications. Different studies have estimated that the proportion of the total citation count contributed by data depositions is around 6–20% [10, 16]. Most of the reanalyses tracked in OmicsDI have been performed using GEO datasets, which might have biased the results to a specific resource. However, our findings show the same patterns in the literature: almost 9,000 datasets have been cited in publications at least once. It is important to highlight that counting direct database citations in the whole text of manuscripts is only possible for open access publications. In the case where the corresponding publications are not open access, dataset identifiers would need to be included in the PubMed abstract to be included in this metric. The coverage of direct citations in publications is therefore limited by this systemic issue. We have found that the transcriptomics community (individual researchers) tend to cite the same datasets more often, with an average of 4 citations per dataset. The most cited dataset is “Transcription profiling of human breast cancer samples - relapse free survival” (E-GEOD-2034), totalling 312 citations. Both metrics, *reanalyses* and *citations*, should be used in combination for a better understanding of the dataset impact. Our results show that both metrics are uncorrelated and should not be aggregated. For example, dataset E-MTAB-513 is among the 10 most cited datasets in the literature, which has been cited 155 times, and reanalysed 4 times. We have decided to only compute and provide these “raw” metrics to the community, rather than combining them into more complex models [32, 33]. However, we have shown that these metrics can be used independently to generate models for classification (**Supplementary Note 4**) [34].

In 2011, Mons *et al.* introduced the idea of *nano-publications*, from where the authors could get credit not only through the actual publication but also through all the knowledge associated with it [7]. In our view, the value of the dataset should not be only associated with the “raw data” or the claims in the publication, but also should be assessed considering all the biological entities supported in knowledge-bases. We have developed the *connections* metric (**Fig. 3d**), which can be used to estimate the impact of a dataset for knowledge-bases, by counting how many biological entities are supported by it.

In addition to the three-metrics used to measure impact, it should be highlighted that OmicsDI routinely uses the number of views/accesses available for each dataset in the resource to rank the results of a search query, using information available from the interaction of other researchers with the indexed datasets. In this context, OmicsDI is monitoring not only the web interface views but also the interaction through the OmicsDI API. In average, every dataset in OmicsDI has been accessed at least 30 times since 2016 (**Supplementary Note 5**). In our view, the number of views/accesses in OmicsDI can be used as a proxy for other relevant metrics such as volume of downloads and number of views in the original data resources. These metrics coming from the original data resources are normally not made publicly available and at present infeasible to retrieve. In fact, at present the first coordinated efforts to gather them in a standard manner are taking place in the context of the ELIXIR framework for European biological data resources [35].

The newly implemented OmicsDI dataset claiming system enables authors, research groups, scientific consortia and research institutions to organize datasets under a unique OmicsDI profile, and for datasets to be added to their own ORCID profiles as well. At the time of writing (March 2018), more than 200 datasets have been claimed into ORCID profiles through OmicsDI. In our view, following the same system for monitoring the impact of individual datasets, these metrics could also be used to measure at least some aspects of the impact of public omics data resources [36, 37]. By March 2018, GEO, PRIDE and MetaboLights datasets are the most cited, re-used, and *connected* in their transcriptomics, proteomics and metabolomics, respectively (**Figure 3**). A common problem of impact evaluation is to compare different fields or topics with the same metrics [38]. Therefore, we recommend the use of these metrics to evaluate datasets within the same omics field, as classified by OmicsDI [1].

## Challenges collecting dataset metrics: recommendations for authors, journals and data resources

Several studies have discussed the challenges of collecting citations for manuscripts and datasets [16, 39]. With the developments of new services such as the EuropePMC API, the compilation of direct citations for datasets has become more feasible [7]. However, in our view, some conventions should be implemented to normalize the way datasets are cited:

i. When the dataset is the main focus of work, dataset identifiers should be used, instead of citing the corresponding publications.
ii. The scientific community needs to develop standard citation strategies for datasets. For example, approximately 60% of the data re-used in one of the drafts of the human proteome papers [39] was collected from public repositories. However, no proper references to the original authors and data are present in the main text of their manuscripts. In order to be able to properly cite the re-use of datasets, new mechanisms should be developed to enable a adequate reporting. OmicsDI has implemented a visualization component (**Supplementary Note 6**) that allows users to cite the corresponding dataset using the FORCE11 Data Citation Synthesis Group (http://www.dcc.ac.uk/resources/how-guides/cite-datasets)[23].
iii. Repositories should make openly available (in an easy to retrieve manner) the links between their reanalysed and the original datasets. Good examples of these links can be found already in Expression Atlas and PeptideAtlas, where every reanalysed dataset references the original ones (**Supplementary Note 7**). Indeed, many databases reference only the associated publications, rather than the actual dataset identifiers. In fact, the correct tracking of datasets in a database by other data resources can help to assess its impact, since it demonstrates that the data they store is actively re-used by (and thus relevant to) the community. Naturally, the same effort should also be made by knowledge bases (e.g. resources including pathways, interactions, gene/protein profiles, etc), to reference the original datasets rather than the publications, in order to recognize that a large part of the biological knowledge is derived from the actual datasets.

We envision that as more and more data is made publicly available, more standardization will be implemented to cross-link resources, manuscripts, datasets and the final biological molecules, making the proposed framework more robust. We expect that any mature omics field should welcome novel insights that can be derived from existing datasets and promote their traceability. We all “stand on the shoulders of giants”. We expect that an improved quantitation of the impact of datasets will help scientists, funders, and research organisations to better value a broader range of “giants” (research products).

## Acknowledgements

YP-R and AZ were supported by US NIH BD2K grant U54 GM114833. GD is supported by EMBL core funding. JAV acknowledges the Wellcome Trust (grant number WT101477MA) and EMBL core funding.

