## Supplementary Materials for "Quantifying the impact of public omics data"

### Supplementary Note 1

Dataset replication is considered a good practice because it guarantees that the dataset will be available even if one of the repositories needs to be closed down or runs out of funding. However, for indexing resources it is desirable to detect that the dataset is the same in the different providers to avoid redundancy. This is a complex task, the identifier of the dataset in the original provider is not always kept in the “replicated” versions. The following dataset (**Figure 1**) was originally deposited in the Gene Expression Omnibus (GEO) ([www.ncbi.nlm.nih.gov/geo/](http://www.ncbi.nlm.nih.gov/geo/)), replicated in the European Nucleotide Archive (ENA) ([www.ebi.ac.uk/ena](http://www.ebi.ac.uk/ena)) and indexed by ArrayExpress ([www.ebi.ac.uk/arrayexpress/](http://www.ebi.ac.uk/arrayexpress/)) and BioProjects ([www.ncbi.nlm.nih.gov/bioproject/](http://www.ncbi.nlm.nih.gov/bioproject/)). All those databases assign different identifiers to the same dataset, GSE59028, SRP044016, E-GEOD-59028 and PRJNA254155, respectively. The current OmicsDI pipeline automatically annotates all the identifiers for a dataset with replicates (**Figure 1**). It also provides a web-interface to manually annotate those datasets that cannot be automatically “merged”.

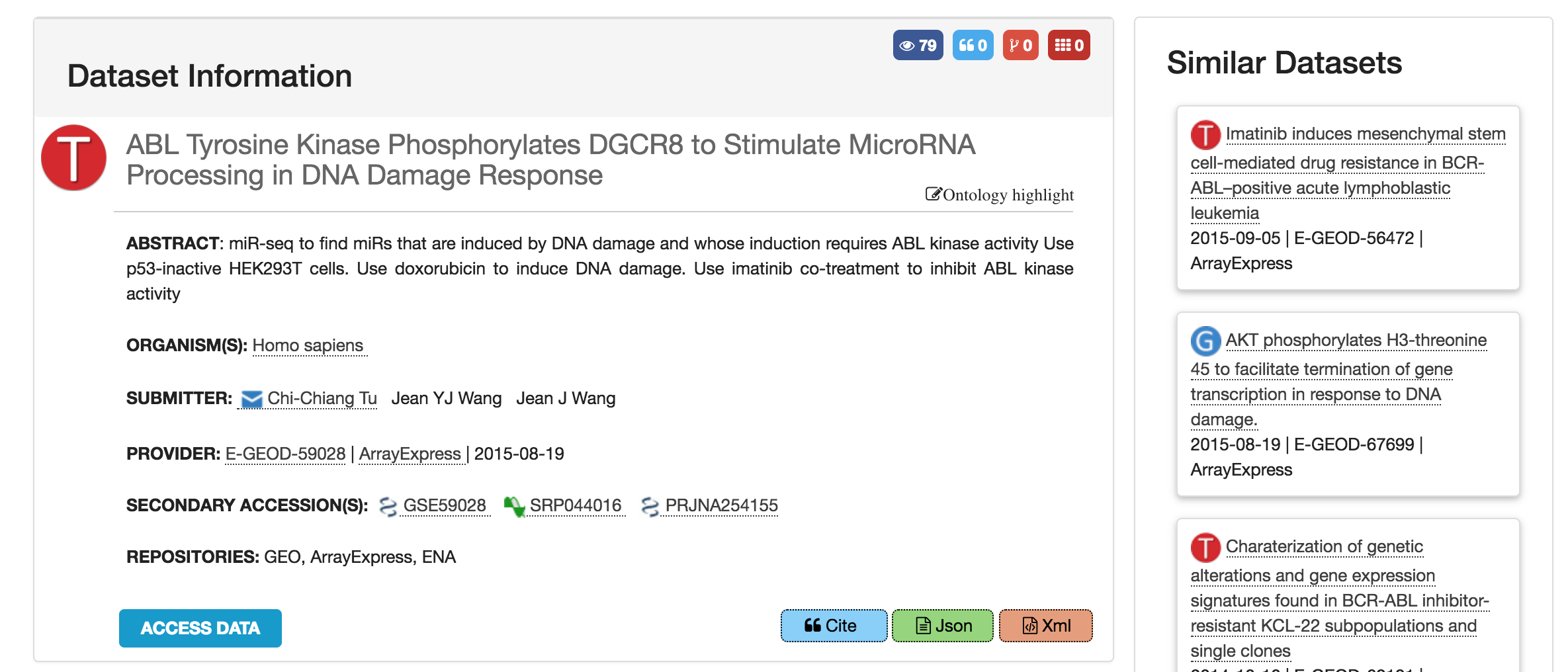

**Figure 1:** ArrayExpress dataset E-GEOD-59028. This dataset can be also found in the original database provider GEO (dataset identifier GSE59028), in the ENA database (identifier SRP044016), and in NCBI BioProjects (identifier PRJNA254155). The OmicsDI canonical identifier pipeline only represents one entry in the OmicsDI resource (in this case the ArrayExpress one) for the corresponding dataset, but also provides all the alternatives/additional identifiers in other data resources.

### Supplementary Note 2

The *connections* metric is calculated by retrieving the references to the dataset from knowledge bases. For example, if a UniProt protein is reported in a proteomics dataset, this counts as a connection to the dataset. The main idea is to keep track of all the biological knowledge generated by one omics dataset. This metric promotes the idea that the value of a dataset is related not only to the number of times it has been downloaded/reused/cited but also how much biological knowledge (e.g. UniProt proteins, Ensembl genes, Interactions or Pathways) is supported by that dataset. **Table 1** shows all the knowledge-base data resources used by OmicsDI to compute the connections to omics datasets. The first column of the table includes the name of the data resource, the second column is the type of data and the last column is the number of entities it contains (by March 2018).

**Table 1**: Knowledge-base data resources included in OmicsDI.

| **Name of the Database** | **Type of Biological Knowledge** | **Number of Biological Entities** |
| --- | --- | --- |
| Assembly | Nucleotide sequences | 144,952 |
| Assembly scaffold (Release) | Nucleotide sequences | 41,759,024 |
| Baseline Expression Atlas Genes | Gene expression | 775,676 |
| ChEMBL Molecule | Bioactive molecules | 1,735,442 |
| ChEMBL Target | Bioactive molecules | 11,538 |
| Coding (Release) | Nucleotide sequences | 356,332,997 |
| DGVa | Genomes & metagenomes | 42,992,846 |
| Differential Expression Atlas Genes | Gene expression | 376,636 |
| Ensembl Gene | Genomes & metagenomes | 3,113,803 |
| Ensembl Genomes Gene | Genomes & metagenomes | 175,370,460 |
| Enzyme Portal | Enzymes | 7,161 |
| HGNC | Genomes & metagenomes | 45,739 |
| IMGT/HLA | Nucleotide sequences | 13,641 |
| IntAct Complexes | Molecular interactions | 2,122 |
| IntAct Interactions | Molecular interactions | 528,593 |
| IntAct Interactors | Molecular interactions | 102,685 |
| IntEnz | Enzymes | 7,686 |
| InterPro | Protein families | 32,568 |
| JPO | Protein sequences | 2,795,178 |
| KIPO | Protein sequences | 518,927 |
| Large assembly | Nucleotide sequences | 44,032,859 |
| Ligands | Bioactive molecules | 19,672 |
| Non-coding (Release) | Nucleotide sequences | 19,045,545 |
| PDBe | Macromolecular structures | 107,958 |
| PfamEntry | Protein families | 16,712 |
| PomBase | Genomes & metagenomes | 6,995 |
| RNAcentral | Nucleotide sequences | 13,353,531 |
| Reactome | Reactions & pathways | 686,345 |
| Rfam | Nucleotide sequences | 2,309,950 |
| Rhea | Reactions & pathways | 43,456 |
| Sequence (Release) | Nucleotide sequences | 205,714,494 |
| TreeFam | Protein families | 15,736 |
| USPTO | Protein sequences | 4,077,368 |
| UniProtKB | Protein sequences | 108,184,003 |
| WormBase ParaSite | Genomes & metagenomes | 2,550,351 |

All these knowledge bases are indexed by the EBI Search resource ([www.ebi.ac.uk/ebisearch/)](http://www.ebi.ac.uk/ebisearch/)) [1]. The EBI search API is used to retrieve all the biological entities that reference the OmicsDI dataset or its corresponding publications (some databases use the publication identifier instead of the dataset accession) (Figure 2a). In addition, we add to the previous list the biological entities that have been reported by the dataset (Figure 2b). By adding the list of referenced biological entities within the dataset we mitigate the time effect between the dataset publication and the use of the data in a knowledge base.

For example, the dataset **E-MTAB-599** has 21,741 connections in EBI SEARCH ([www.ebi.ac.uk/ebisearch/search.ebi?db=allebi&query=E-MTAB-599&requestFrom=searchBox)](http://www.ebi.ac.uk/ebisearch/search.ebi?db=allebi&query=E-MTAB-599&requestFrom=searchBox)); 115 Nucleotide sequences, 21,572 Gene expression profiles, 54 Samples & ontologies mentions (connection set **A**). Then, we query the EBI Search for the manuscript associated with the dataset (<https://www.ebi.ac.uk/ebisearch/crossrefsearch.ebi?db=europepmc&id=21921910&ref=genomes&requestFrom=relatedData-xref)>. The results showed 1,689,177 new connections from the Database from Genomics Variation Archive (connection set **B**). In addition, the 21,572 reported in the dataset considered (connection set **C**). The final connection metric number is the union of the three sets (**A** U **B** U **C**). By using the union of the three groups we avoid duplications of connections among providers.

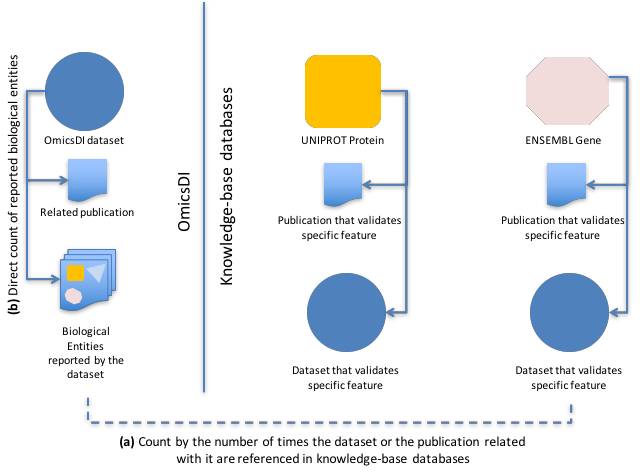

**Figure 2**: Schema of the reference system used in OmicsDI based on the EBI Search architecture. Each OmicsDI dataset is supported by a publication and a list of biological entities in knowledge bases such as UniProt, Ensembl, etc. In the same way, entities in knowledge bases are supported by publications or datasets. (**a**) The connections pipeline in OmicsDI counts connections to a dataset or its related publication if referenced in a knowledge base resource. In addition, (**b**) the pipeline uses the list of biological entities reported by the dataset to complement the previous lists in case the dataset hasn’t been added to the knowledge base. The final connection metric is the **union** of all the collected lists.

### Supplementary Note 3

OmicsDI has implemented a dataset claiming system that allows researchers to get credit for generating and sharing their datasets. The platform is built on top of three main components: i) a log-in system based on ORCID (**Figure 3a**); ii) a dataset search-based claiming system that allows users to search and add the relevant datasets to their OmicsDI profiles (Figure 3**b**); and iii) a user profile enables users to update datasets in their OmicsDI profile (**Figure 4a**), and synchronize them with ORCID (**Figure 4a**). When the profile is synchronized with ORCID, the user’s datasets are added to their ORCID profile (**Figure 4b**). The claiming step is based on the *principle of trust* as in other popular analogous resources for publications such as Google Scholar or ResearchGate.

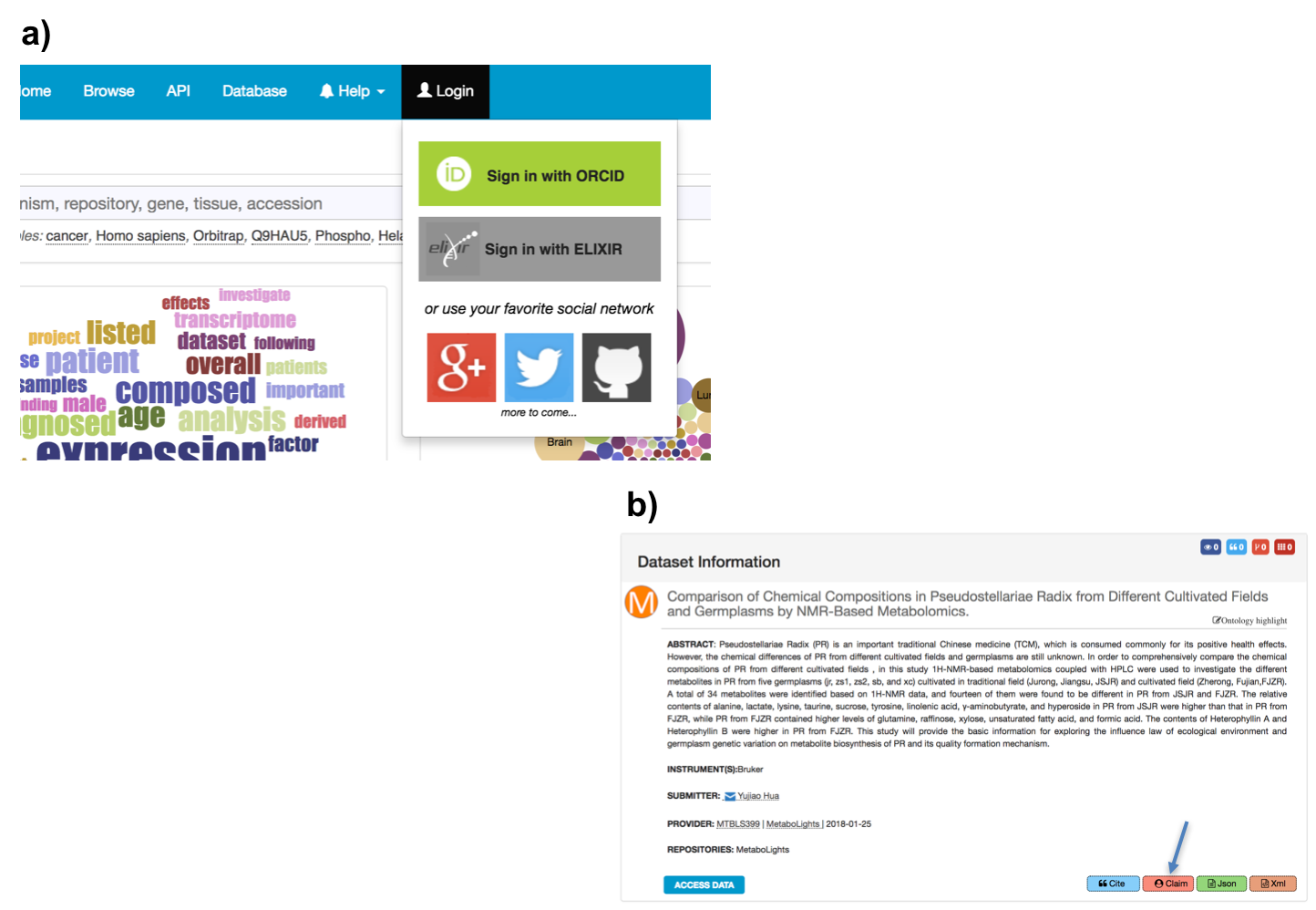

**Figure 3**: **(a)** OmicsDI has provided a simple log-in and personal profile by using the ORCID authentication system. **(b)** When the researchers are logged in, a new option appears at the bottom of each dataset page, which allows them to claim the corresponding dataset.

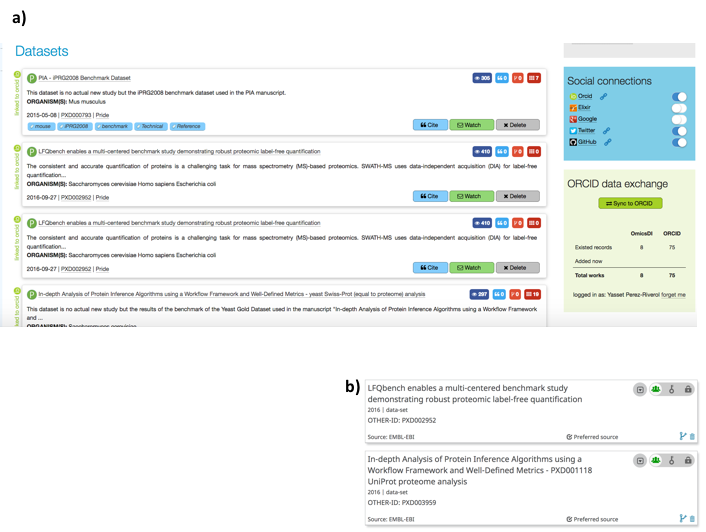

**Figure 4**: (**a**) Datasets claimed by a specific user are listed in their OmicsDI profile. OmicsDI enables users to update their profile. A profile can be made public and the URL can be shared in public domain, to demonstrate the impact of the author’s datasets. **(b)** Each researcher’s dataset list can be synchronized with their own ORCID profile.

### Supplementary Note 4

We have combined the data *citation* and *reanalyses* metrics using a simple *k-means* clustering algorithm to identify which are the datasets that are highly-cited, highly-reanalysed, moderately-cited, and moderately-reanalysed (**Figure 5**). This simple classification enables users and services to search and find datasets in a resource such as OmicsDI, with contains more than 100,000 datasets. The current results show that there are 113 (highly-cited), 682 (moderately-cited), 245 (moderately-reanalysed), 8 (highly-reanalysed) datasets.

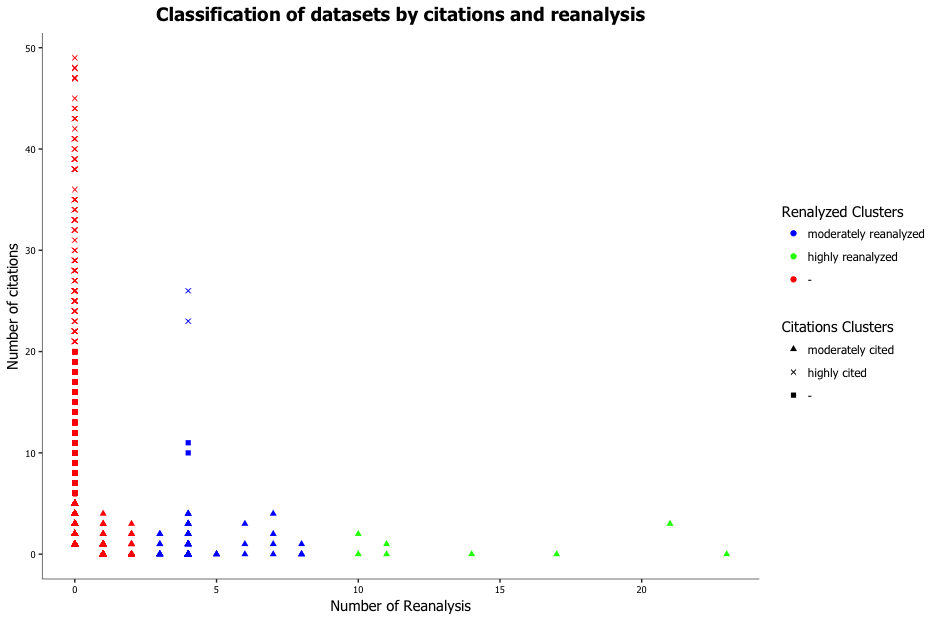

**Figure 5**: Classification of datasets into four different categories (highly-cited, highly-reanalysed, moderately-cited, moderately-reanalysed) using the *k-means* cluster algorithm.

### Supplementary Note 5

| Omics Type | Number of Views | Avg. Number Views |
| --- | --- | --- |
| Transcriptomics | 3,727,514 | 29.9 |
| Proteomics | 490,805 | 52.1 |
| Metabolomics | 38,686 | 38.0 |
| Genomics | 228,654 | 31.3 |
| Biological models | 167,405 | 19.9 |

**Table 2**: Average number of views/access for datasets in OmicsDI, split by omics type.

### Supplementary Note 6

Data citation is a crucial component of OmicsDI. We have implemented a simple visualization component that allows users to export dataset citations using two citation styles (APA and AMA) (**Figure 6**).

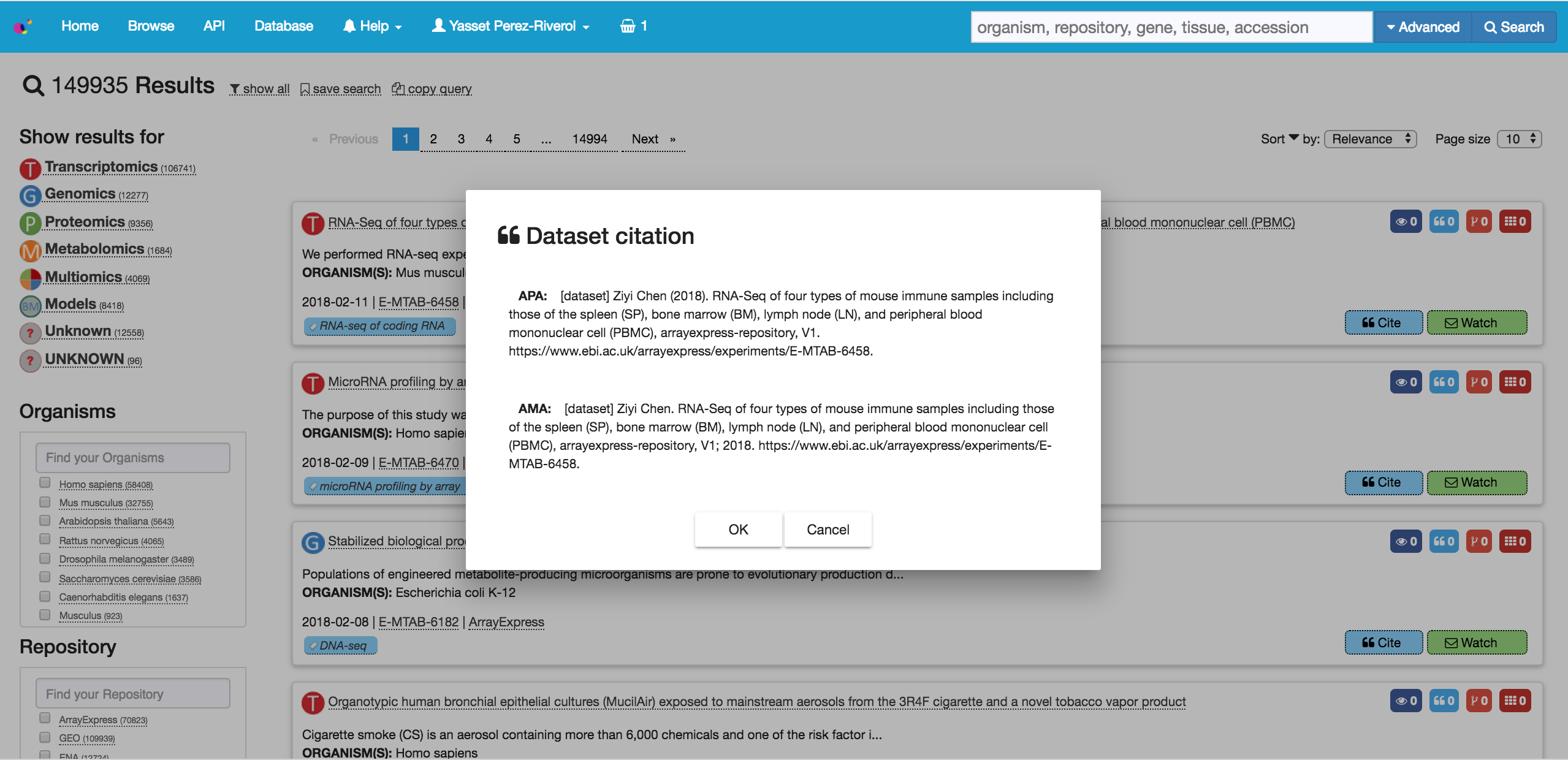

**Figure 6**: Dataset citations in OmicsDI can be exported in APA and AMA styles.

### Supplementary Note 7

The Expression Atlas database provides for each dataset (reanalysis) the reference to the original dataset in ArrayExpress (**Figure 7**, see bottom of the page – Resources section). The reanalysis reference to the original dataset allows OmicsDI to monitor the number of times a dataset in one of the Archive (original providers) is used by another database or resource.

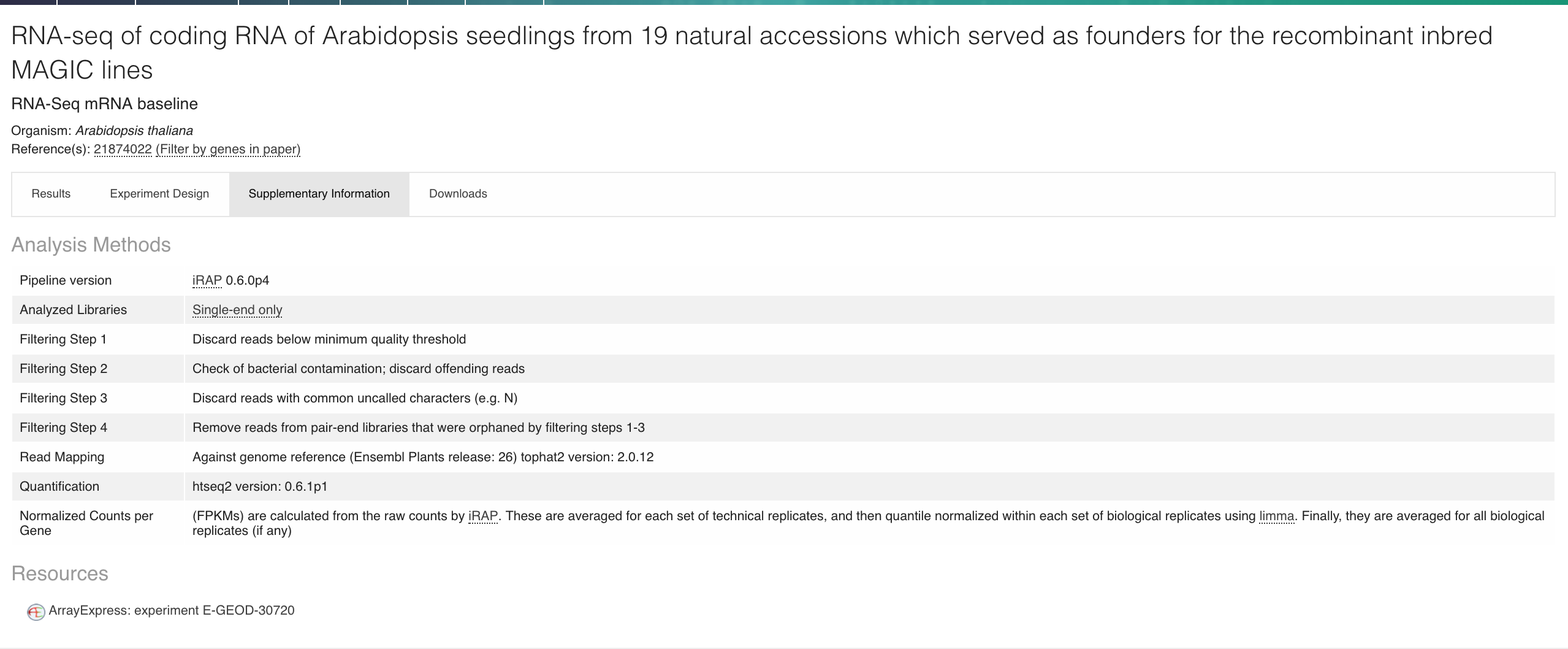

**Figure 7**: Expression Atlas’ direct reference to the original re-used transcriptomics dataset originally deposited in ArrayExpress.
